# Early transcriptional landscapes of *Chlamydia trachomatis*-infected epithelial cells at single cell resolution

**DOI:** 10.1101/724641

**Authors:** Regan J. Hayward, James W. Marsh, Michael S. Humphrys, Wilhelmina M. Huston, Garry S.A. Myers

## Abstract

*Chlamydia* are Gram-negative obligate intracellular bacterial pathogens responsible for a variety of disease in humans and animals worldwide. *C. trachomatis* causes trachoma (infectious blindness) in disadvantaged populations, and is the most common bacterial sexually transmitted infection in humans, causing reproductive tract disease. Antibiotic therapy successfully treats diagnosed chlamydial infections, however asymptomatic infections are common. High-throughput transcriptomic approaches have explored chlamydial gene expression and infected host cell gene expression. However, these were performed on large cell populations, averaging gene expression profiles across all cells sampled and potentially obscuring biologically relevant subsets of cells. We generated a pilot dataset, applying single cell RNA-Seq (scRNA-Seq) to *C. trachomatis* infected and mock-infected epithelial cells to assess the utility of single cell approaches to identify early host cell biomarkers of chlamydial infection. 264 time-matched *C. trachomatis*-infected and mock-infected HEp-2 cells were collected and subjected to scRNA-Seq. After quality control, 200 cells were retained for analysis. Two distinct clusters distinguished 3-hour cells from 6- and 12-hours. Pseudotime analysis identified a possible infection-specific cellular trajectory for *Chlamydia*-infected cells, while differential expression analyses found temporal expression of metallothioneins and genes involved with cell cycle regulation, innate immune responses, cytoskeletal components, lipid biosynthesis and cellular stress. Changes to the host cell transcriptome at early times of *C. trachomatis* infection are readily discernible by scRNA-Seq, supporting the utility of single cell approaches to identify host cell biomarkers of chlamydial infection, and to further deconvolute the complex host response to infection.

## Introduction

*Chlamydia* are Gram-negative obligate intracellular bacterial pathogens that cause disease in humans and a wide variety of animals. In humans, *Chlamydia trachomatis* typically infects cells within the ocular and genital mucosa, causing the most prevalent bacterial sexually transmitted infections (STI) [1], inducing acute and chronic reproductive tract diseases that impact all socioeconomic groups, and trachoma (infectious blindness) in disadvantaged populations [2]. Disease outcomes arise from complex inflammatory cascades and immune-mediated host processes that can lead to tissue damage and fibrotic scarring in the upper genital tract or the conjunctiva [3,4]. Reproductive tract disease outcomes include pelvic inflammatory disease (PID), preterm delivery, ectopic pregnancy, hydrosalpinx, tubal factor infertility (TFI) and chronic pelvic pain in women, as well as epididymitis, testicular pain and infertility in men. Antibiotic therapy with azithromycin or doxycycline successfully treats diagnosed infections, however asymptomatic infections are common [5,6], and so treatment cannot be administered. Asymptomatic infection rates are estimated to exceed diagnosed infection rates by at least 4.3-fold [5].

*Chlamydia* have a unique biphasic developmental cycle with distinct morphological forms. The cycle begins with attachment and entry of the infectious elementary bodies (EBs) into host cells, typically mucosal epithelial cells. After entry, EBs reside within membrane-bound vacuoles that escape phagolysomal fusion [7]. Differentiation into the replicating reticulate bodies (RBs) occurs within the first 2-3 hours, followed by continued growth of the inclusion accommodating the increased number of RBs. Over the course of infection, *Chlamydia* parasitises and modifies the host cell by deploying type III effectors and other secreted proteins [8], which also facilitate invasion, internalisation, and replication, while countering host cell defences [9,10]. At the end of the developmental cycle, RBs asynchronously transition back into EBs (~20-44 hours) and, through either extrusion or host cell lysis (~48-70 hours), are released to repeat the cycle [11]. Chlamydial transcriptomes have been examined over the developmental cycle, in EBs and RBs, and in different chlamydial species [12–15]. Dual RNA-Seq [16] has allowed the transcriptomes of both *C. trachomatis* and infected epithelial cells to be profiled simultaneously, identifying previously unrecognised early chlamydial gene expression and complex host cell responses [17]. However, the derived transcriptional profiles represent averaged gene expression over the population of cells sampled [18]. Subsets of cells with dominant gene expression profiles can skew the analysis [19], possibly obscuring other potentially important cell subsets and their transcriptional profiles [20,21]. By examining the expression profiles of individual cells, single cell RNA sequencing (scRNA-Seq) can minimise these biases, enabling a deeper understanding of population heterogeneity, cell states and interactions, and gene regulation [22,23]. scRNA-Seq and other single cell methods have been instrumental in discovering new cell types [22] and advancing the understanding of many disease states [24], particularly tumour heterogeneity [25,26], hematopoiesis [27] and embryonic development [28]. Applications of scRNA-Seq to pathogen-infected cells are more limited so far, but are exemplified by studies that show the heterogeneity of macrophage responses to *Salmonella enterica* serovar Typhimurium infection [29], the high degree of cell-cell transcriptional variation induced by influenza virus infection [30], and the characterization of lymph node-derived innate responses to bacterial, helminth and fungal pathogens [31].

Here we explore the application of single cell analysis methodologies to *Chlamydia*-infected cells, with the goals of identifying host cell developmental-stage biomarkers, and to assess the utility of these methodologies for deciphering chlamydial biology in cells and tissues. We generated a pilot scRNA-Seq dataset of time-matched infected and mock-infected HEp-2 epithelial cells *in vitro* encompassing the early chlamydial developmental cycle (3, 6 and 12 hours). We show that infection responsive changes to the early host cell transcriptome are readily discernible by scRNA-Seq, supporting the potential for host derived infection biomarkers.

## Results

### Quality assessment and filtering

Raw sequencing reads were demultiplexed using DeML [32], yielding 1.03 billion sequence reads (**Supplemental File 1**). Single cell datasets were removed from subsequent analyses if they contained less than 1 million reads after trimming and alignment, and less than 5,000 counted features (genes). Further quality assessment measures ensured that sequence reads mapped across all chromosomes and that the majority of reads mapped to protein-coding genes (**Supplemental File 2**). Single cell datasets were pooled and subjected to additional quality assessment steps, including examining rRNA as a measure of depletion success and mitochondrial gene expression as an indicator of cell stress [33], as both are potential sources of bias (**Supplemental File 3**). During quality control, the mock-infected cells at 3 hours failed to pass cut-offs, and were excluded from further downstream analysis (**Supplemental File 3**). After all quality measures, 200 high quality single cells remained across the three times.

### Removal of confounding effects

To normalise by library size, Scran’s single-cell specific method was used to deconvolute library size factors from cell clusters [34]. We applied RUVSeq [35] to identify and remove further confounding effects, including differences between batches of sequenced cells. Reduction of variation was confirmed in relative log expression (RLE) plots (**Supplemental File 4A**). Density curve plots further show the effect of removing variability from the raw counts, after library size normalisation, and after removing further confounding effects (**Supplemental File 4B**). The PCA bi-plot (**Supplemental File 4C**) shows the structure of the data and grouping of the cells following these steps. By examining the underlying variables driving PC1 variation, we found that total read counts and time post infection account for 99% of the total variation (**Supplemental File 4C**), confirming that most variation is not from experimental factors. In addition, doublets (where at least two cells are contained in a droplet) can skew the resulting expression profiles, adding a further confounding factor. The C1 platform has been associated with a high doublet rate [36]. Due to this high reported rate, we ran different tools to identify doublets, confirming that our data had minimal detected doublets (**Supplemental File 5**).

### Cell cycle classification

Due to the constraints imposed by chlamydial infection within *in vitro* tissue culture and, given the potential for cell-cell variability despite infection synchronisation, we expected to observe a range of cell cycle stages in our data (**Supplemental File 6**). Two of the three stages (G1 and G2/M) show more than double the number of cells from 3 to 6 hours, while DNA synthesis (S) is the only cell cycle stage with a decrease in the number of cells from 3 to 12 hours. However, despite these trends, no distinct cell cycle clusters are apparent (**Supplemental File 6B)**. In addition, there is no clustering between cell cycle state and time post-infection, or infection condition (infected *vs* mock-infected). Although we identify cell cycle stage as a likely confounding effect [37] that was removed from our subsequent analyses, it may be relevant to the infection and growth strategies of *Chlamydia*. For example, while infected cells can still grow and divide, the burden of infection causes these cells to proliferate more slowly than uninfected cells, resulting in dividing cells which may be more or less susceptible to infection [38]. Additionally, chlamydial infectivity has been related to distinct cell cycle phases, where infection can modulate cell cycle parameters [39].

### Clustering demonstrates transcriptional heterogeneity of infected epithelial cells over the early chlamydial developmental cycle

Unsupervised clustering identified two distinct clusters across the three time points (**Figure 1C** and **Supplemental File 7**). Cluster 1 contains only 3-hour infected cells, while cluster 2 contains a mixture of cells from 6 and 12 hours, with no clear separation between infected and uninfected conditions. We used k-nearest neighbour smoothing (kNN-smoothing) to further reduce scRNA-Seq-specific noise [40] within the expression matrix. The resulting PCA plot recapitulated the clusters identified above; indicating that the previous clustering result was not influenced by noise-related factors. Additional clustering analyses were performed to identify any sub-populations within each cluster on the basis of experimental factors such as time or infection status (**Supplemental File 8**). *Chlamydia*-infected cells again clustered into two main groups, closely matching the overall clustering that separated the 3 hours cells from 6 and 12 hours, with no further sub-clustering evident.

**Figure 1:**
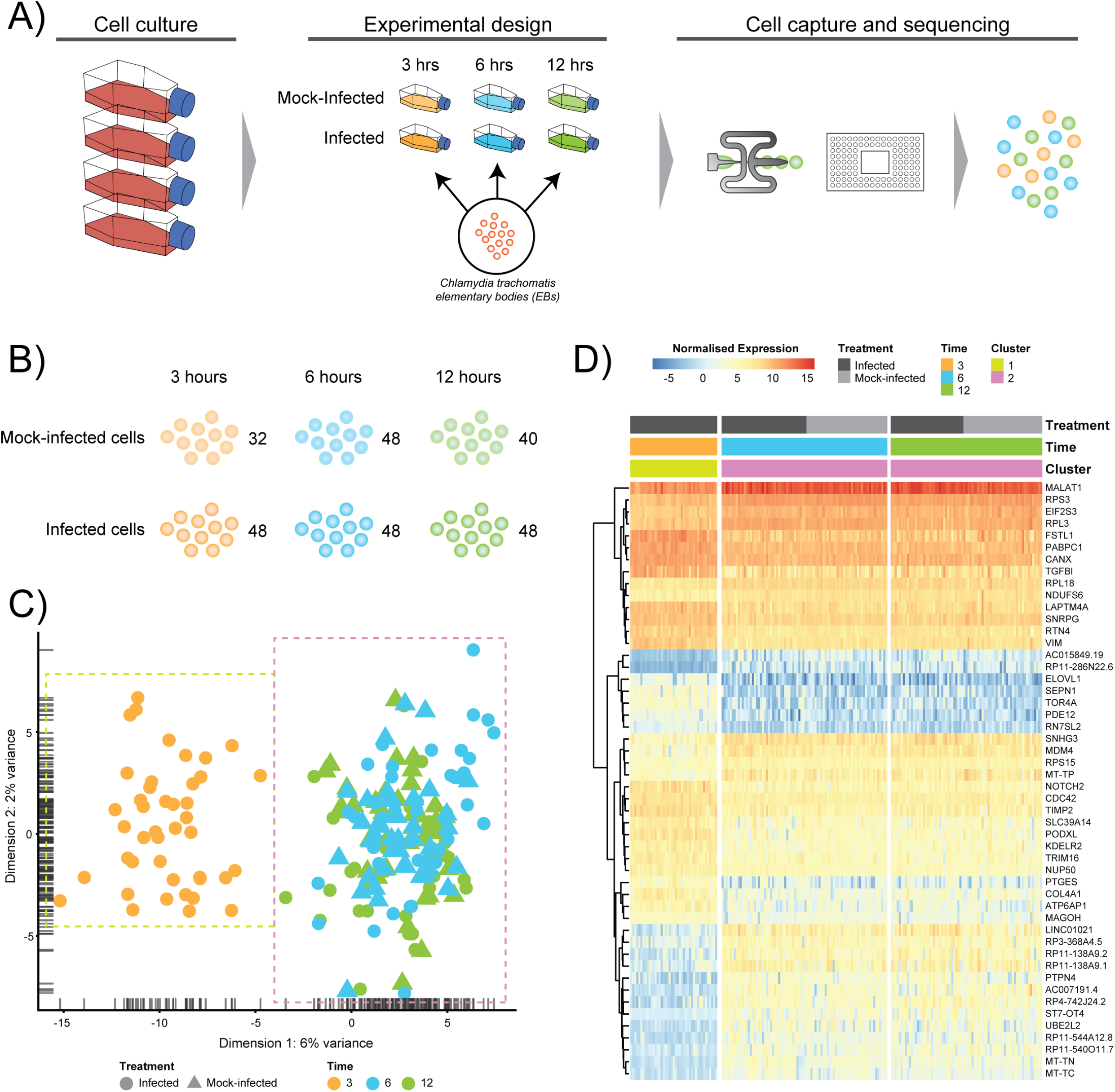
Experimental design and analysis. **A)** Experimental design of *C. trachomatis* E and time-matched mock-infections using HEp2 epithelial cell monolayers at 3, 6 and 12 hours post-treatment, prior to capture and scRNA-Seq library preparation on the Fluidigm C1 instrument. **B)** Numbers of captured and sequenced single cells by experimental condition and time. **C)** Unsupervised clustering identifies two primary clusters. Cluster 1 contains all 3-hour cells, while cluster 2 contains all 6− and 12-hour cells. **D)** Putative marker genes identified by hierarchical clustering.

### Pseudotime analysis over the early chlamydial developmental cycle

Clustering analyses demonstrate that both infected and uninfected cells have minimal cluster separation at 6 and 12 hours **(Figure 1C)**. We applied pseudotime analysis to further deconvolute cellular trajectories that may follow a time course or biological mechanisms such as differentiation or infection [41,42]. Pseudotime analysis of *Chlamydia*-infected cells alone predicted 3 distinct cell states (**Supplemental File 9A**). Cell state 1 contained 3-hour cells, state 2 contained a mixture of 3, 6− and 12-hour cells, and state 3 contains a mixture of 6 and 12 hour cells (**Supplemental File 9B**). The line connecting the 3 cell states (minimum spanning tree) does not provide a realistic linear trajectory of the infection course from 3-12 hours. Manually increasing the number of states does provide a more realistic trajectory (**Supplemental File 9C**); however, the similarity of 6− and 12-hour cells (both mock-infected and infected states) will require more cells to accurately capture any putative sub-stages of infection.

We used cell state prediction to identify any shared characteristics of infected cells with mock-infected cells, which could classify cells that were either not infected or were perhaps able to overcome the bacterial infection early. This analysis identified only two cell states (**Supplemental File 9CA**), again recapitulating the clustering results. By manually increasing the number of cell states to 7, smaller sub-clusters within the two main clusters became evident (**Supplemental File 9C**). However, we still observe a mixture of infected and mock-infected cells within each sub-cluster (albeit with small numbers of cells), further highlighting the transcriptional similarity between infected cells at 6 and 12 hours.

### Differentially expressed genes in *Chlamydia*-infected and mock-infected cells highlight infection mechanisms

Subsets of genes with significant expression differences between the two primary clusters were examined in order to identify putative host cell marker genes that can distinguish different times post-infection (**Figure 1D**). At 6 and 12 hours, genes associated with processes governing RNA and protein metabolism (*RPS3*, *RPL3*, *RPS15*, *RPL18*, *PABPC1*, *MAGOH* and *EIF2S3*) predominate, with most showing increased expression. Increased expression of vimentin (*VIM*), a type III intermediate filament (IF) present in the cytoskeleton and involved in maintaining cell shape and integrity [43], distinguishes the 3 hour cluster.

We further examined differentially expressed (DE) genes firstly by comparing infected and mock-infected cells at 6 and 12 hours respectively (cluster 2), and secondly by comparing the 3 hour infected cells (cluster 1) against cluster 2, as the 3 hour mock-infected cells were removed after initial quality control steps. At 6 hours, 44 DE genes were identified (13 up-regulated and 21 down-regulated) (**Supplemental File 10A**), including three up-regulated metallothionein (MT) genes (*MT1E*, *MT2A* and *MT1X*). MT up-regulation occurs in response to intracellular zinc concentration increases, reactive oxygen species (ROS) and proinflammatory cytokines [44]. Intracellular zinc concentrations are an integral component of immunity and inflammation, and zinc deficiency results in an increased susceptibility to infection [45]. MTs may also have a role in protecting against DNA damage and in apoptosis, as well as regulating gene expression during the cell cycle [46], which are likely to be relevant at 6 hours post infection. Down-regulated pathways at 6 hours were dominated by three genes *HSP90AA1* (Heat Shock Protein 90 Alpha Family Class A Member 1), *TUBB* (Tubulin Beta Class I), and *TUBA4A* (Tubulin Alpha 4a), which are linked to the cell cycle, specifically centrosome maturation and microtubule assembly mediating mitosis (**Supplemental File 10B**). *Chlamydia*-infected cells are still able to undergo mitosis, however mitosis-related defects do occur during chlamydial infection. These include an increase in supernumerary centrosomes, abnormal spindle poles, and chromosomal segregation defects, and result in a heavily burdened cell [47,48]. Cells that recover from infection are still likely to contain chromosome instabilities, which can then be passed down to uninfected daughter cells [48].

At 12 hours, there is an increase in DE genes (245) with 98 up-regulated and 147 down-regulated (**Supplemental File 10A**). We continue to see up-regulated genes that are likely part of a continued immune response to infection, including two MTs (*MT1M* and *MT1E*), *TRIM25* (Tripartite Motif Containing 25), *ISG15* (*ISG15* Ubiquitin Like Modifier), *HLA-A* (Major Histocompatibility Complex, Class II, DR Beta 1), *IFIT3* (Interferon Induced Protein With Tetratricopeptide Repeats 3), *OASL* (2’-5’-Oligoadenylate Synthetase Like), *IL6* (Interleukin 6), and genes associated with cholesterol and fatty acid synthesis (**Supplemental File 10B**). The exploitation of a variety of host lipids by *Chlamydia* to subvert intracellular signalling, survival and growth is well established [49–51]. All down-regulated pathways at 12 hours indicate that *Chlamydia*-infected cells are exhibiting stress responses. DNA damage as part of the cell cycle, and repair pathways are enriched, possibly representing a continuation of infection stresses at 6 hours, and likely indicative of further *Chlamydia*-induced interruption of the cell cycle. Notably, two p53 associated pathways were enriched from associated genes. p53 expression tightly controls the cell cycle and is modulated in response to activities including cell stress, DNA damage, as well as bacterial infection [52]. *Chlamydia-*induced down-regulation of p53 may help to protect infected cells against death-inducing host responses, thus allowing chlamydial survival [53]. Only four DE genes from 6 and 12 hours overlap. The two up-regulated genes were *DUSP5* (Dual Specificity Phosphatase 5) and *MT1E* (metallothionein 1E), while the two down-regulated genes were *TUBA4A* and *HSP90AA1*.

Comparing cluster 1 (3-hour infected cells) against cluster 2 (DE genes from 6 and 12 hours) demonstrates a substantial number of DE genes (2,291 up and 3,487 down-regulated) (**Supplemental File 10A**). The down-regulated pathways have low combined scores compared to the up-regulated pathways, which may be explained by the large number of down-regulated non-coding RNAs (ncRNAs), which are typically not incorporated into the underlying enrichment analyses. Three of the up-regulated pathways are associated with infection (*Infectious disease*, *Influenza life cycle* and *Influenza infection*) (**Supplemental File 10B**), demonstrating that general infection mechanisms are the key differences between these temporally defined clusters, and further confirming that the majority of cells were successfully infected.

## Discussion

To better understand bacterial pathogenesis and resulting disease outcomes, it is critical to understand functional changes to specific cell populations of infected and neighbouring cells, and recruited immune cells in the infected tissue context. This is especially relevant for *Chlamydia* which, due to its obligate intracellular niche and distinct morphologies, has long been refractory to research. As a result, many infection and disease processes at the cellular and tissue level remain largely unknown or poorly characterised *in vivo*. Gene expression profiling and other genome-scale analyses are powerful techniques for deconvoluting these interactions and processes. However, these genome-scale analyses have typically been performed on bulk cell populations, i.e.; infected cell monolayers *in vitro*, or selectively sorted/purified subsets of cell populations. Such bulk cell approaches can miss cell-cell variability, or cells that contribute to overlapping phenotypic characteristics, potentially masking critical biological heterogeneity as irrelevant signals from non-participating cells that can skew the average. This may influence the understanding of multifactorial and dynamic processes, such as inflammation and fibrosis during ascending chlamydial infection.

We describe the first application of scRNA-Seq to *Chlamydia*-infected cells. This pilot dataset, comprising 264 single infected and mock-infected cells encompassing three early times of the *in vitro* chlamydial developmental cycle, was designed to examine the feasibility of using single cell approaches to identify host-derived transcriptional biomarkers of chlamydial infection. After quality assessment and filtering measures, we retained 200 high quality, *C. trachomatis*-infected and mock-infected cells.

Clustering, pseudotime and cell state prediction analyses confirmed that these cells were infected, and demonstrated that infected cells at 3 hours are readily distinguishable from infected and mock-infected cells at 6 and 12 hours. Curiously, cells at 6 and 12 hours clustered together and could not be further deconvoluted from each other, showing that host cell transcription at these times is broadly similar. A recent FAIRE-Seq analysis of *Chlamydia*-infected epithelial cells, examining patterns of host cell chromatin accessibility over the developmental cycle [54], found that 12 hours post infection was relatively quiescent in terms of host cell transcriptional activity. This finding is reflected by our scRNA-Seq analyses here and may be extended to 6 hours post-infection. In addition, both *Chlamydia*-infected and mock-infected cells at 6 and 12 hours clustered together. One interpretation of this phenomenon is that these early infection times represent a period where the ongoing establishment of the inclusion and chlamydial division after initial entry and infection events is largely cryptic to the host cell as manifested by transcriptional processes.

Differential expression comparing cells between clusters identified both conserved and temporally-specific gene expression over the times examined. Many of the identified pathways and genes are relevant to chlamydial infection processes, including metallothioneins, innate immune processes, cytoskeletal components, lipid biosynthesis and cellular stress. In addition, genes and pathways related to the cell cycle are down-regulated; while this could be an off-target effect of infection that does not benefit *Chlamydia*, interference with the cell cycle may constitute an infection strategy as *in vivo* cells will be at different cell cycle stages, and thus some may be more or less susceptible to infection.

These pilot analyses demonstrate that distinct host cell transcriptional responses to infection are readily discernible by single cell approaches even at the early stages of the chlamydial developmental cycle, yielding robust data and confirming that host cell-derived transcriptional biomarkers of chlamydial infection are identifiable. Single cell genome-scale approaches applied to *Chlamydia*-infected and neighbouring cells, recruited immune cells from inflammatory processes, and structural cells obtained from clinical swabs or *ex vivo* tissues, are likely to lend significant insight to the complex processes that underpin chlamydial infection and the associated inflammatory disease outcomes.

## Methods

### Cell culture and infection

HEp-2 cells (American Type Culture Collection, ATCC No. CCL-23) were grown as monolayers in 6 x 100mm TC dishes until 90% confluent. Monolayers were infected with *C. trachomatis* serovar E in SPG as previously described [55]. Additional monolayers were mock-infected with SPG only. The infection was allowed to proceed 48 hours prior to EB harvest, as previously described [55]. *C. trachomatis* EBs and mock-infected cell lysates were subsequently used to infect fresh HEp-2 monolayers. Fresh monolayers were infected with *C. trachomatis* serovar E in 3.5 mL SPG buffer for an MOI ~ 1 as previously described [55], using centrifugation to synchronize infections. Infections and subsequent culture were performed in the absence of cycloheximide or DEAE dextran. A matching number of HEp-2 monolayers were also mock-infected and synchronised as above using uninfected cell lysates. Each treatment was incubated at 25°C for 2h and subsequently washed twice with SPG to remove dead or non-viable EBs. 10 mL fresh medium (DMEM + 10% FBS, 25 μg/ml gentamycin, 1.25 μg/ml Fungizone) was added and cell monolayers incubated at 37°C with 5% CO_**2**_. Three biological replicates of infected and mock-infected dishes per time were harvested post-infection into single cell populations by trypsin in sterile PBS prior to immediate single cell capture and library preparation. We note that the experimental design employed here will not distinguish *Chlamydia*-mediated effects from infection-specific or non-specific epithelial cell responses. We further note that these *in vitro* infections are synchronized by design, thereby minimising the degree of heterogeneity within each infection time in order to enable more accurate examination of temporal effects.

### Library preparation and sequencing

A Fluidigm C1 instrument was used for cell capture and subsequent library preparation. The resulting 264 single cell libraries were sequenced using the Illumina HiSeq 4000 platform (150bp paired-end reads) across three batches. Each plate was designed with a balanced distribution of time points and conditions.

### Pre-processing and quality control

Raw sequencing reads were demultiplexed using DeML [32] with default settings. Trim Galore (v.0.4.3) [56] was used to trim adaptors and low quality reads. Confirmation of the removal of adaptors, low quality reads and other quality control measurements was performed with FastQC (v.0.11.5) [57]. Reads were subsequently aligned to the human genome version (GRCh 38.87) with STAR (v.2.5.1a) [58] retaining paired and unpaired mapped reads that were merged in to a single BAM file.

FastQ-Screen (v.0.11.1) [59] was used to screen for sources of contamination across all cells. This output and low mapping rates confirmed the removal of all 3-hour uninfected cells, due to extremely low mapping rates to the Human genome and high mapping rates to other organisms. Features of the remaining cells were counted with FeatureCounts (v.1.5.0-p3) [60]. MultiQC (v.1.0) [61] was used throughout the previous steps, combining output from each piece of software to easily make comparisons between batches and across time points.

### Identifying outlier cells based on filtering

Counted features were imported into Scater (v.1.5.11) [62], where subsequent quality control reduced the total number of cells from 264 to 200. The filter settings were comprised of four steps: 1) total mapped reads should be greater than 1,000,000; 2) total features greater than 5,000; 3) expression from mitochondrial genes less than 20% of total expression; and 4) expression from rRNA genes comprise less than 10%.

### Removing confounding effects

Cell cycle classification was performed using Cyclone [63] prior to filtering out low abundance genes, as recommended. To account for the differences in library sizes between cells, the deconvolution method from Scran (v.1.6.0) [63] was used. Further confounding effects such as cell cycle and sequencing batch effects were removed using the RUVs method of RUVSeq (v.1.12.0) [35] using k=4. Doublet detection was performed using Scrublet (v.0.1) [64] and DoubletDetection (v.2.4) [65].

### Clustering

Unsupervised clustering was performed using the Single-Cell Consensus Clustering (SC3) package (v.1.10.1) [66]. Two clusters (k) were chosen based on automatic prediction by SC3 after iterating through a range of k (2:10). Higher values of K were examined; however, two clusters remained the best fit to this data as assessed by various internal plots including consensus matrices, silhouette plots and cluster stability plots. To further confirm the two clusters, the KNN-Smoothing [40] (K-nearest neighbour smoothing) function (v.1) was also applied to different transformations of the library normalised data.

### Pseudotime analysis

TSCAN (v.1.16.0) [42] was used to perform pseudotime analysis. When all cells were analysed, the “pseudocount” and “minexpr_value” flags were set to 0.5 in pre-processing to allow more features to be selected, resulting in an increase of cells with assigned cell states, especially when the number of states was manually increased. The default pre-processing settings were used to examine infected cells alone.

### Differential expression

Most scRNA-Seq differential expression software only allows for direct comparisons, such as cluster comparisons. As our experimental design examined both infected and mock-infected cells, we used edgeR (v.3.24.3) [67], as it provides better functionality for more complex comparisons. In addition to including RUVSeq factors of unwanted variation to the model matrix, the dispersion trend was estimated using “locfit” and “mixed.df” flags set to true. Resulting p-values were adjusted using a false discovery rate (FDR) < 0.05; significant genes were examined with Reactome [68].

## Availability of supporting data and materials

The data set supporting the results of this article is available in the GEO repository, [GSE132525].

## Additional Figure Legends

**Supplemental File 1. Demuxing results comparing Illumina and DeML**

Demuxing comparison between the standard Illumina software and DeML. 1.8-5.5% additional reads were recovered over each cell batch.

**Supplemental File 2. Biotype distribution**

**A)** Distribution of different gene biotypes by time. **B)** Total gene expression by biotype. **C)** Total gene expression by chromosome location.

**Supplemental File 3. Contamination of 3-hour mock-infected cells**

Examples of two 3-hour mock-infected cells with unusual mapping to 12 different genomes, indicating cross-contamination. All 3-hour mock-infected cells showed similar profiles and were removed from further downstream analysis.

**Supplemental File 4. Identifying and removing confounding effects**

**A)** Relative log expression (RLE) plot of gene expression from all cells after removal of confounding effects using RUVSeq. **B)** Density curve distribution showing variability associated to key variables from raw counts, after library normalisation, and after using RUVSeq respectively. **C)** PCA plot demonstrating two-dimensional structure of cell expression profiles after removing confounding effects. Two variables (Total features and Time-point) account for 99% of the variability at component 1 (PC1).

**Supplemental File 5. Doublet detection**

**A)** Scrublet identifies two cells as possible doublets, red bars. **B)** DoubletDetector found no cells exhibited doublet characteristics.

**Supplemental File 6. Cell cycle classification**

**A)** Cell cycle classification of single cells after removing outliers. **B)** PCA plot examining cell-cycle related trends by time-point and infection status.

**Supplemental File 7. Clustering**

**A)** SC3 consensus matrix predicted 2 clusters, dark red colouring. **B)** Silhouette plot of the consensus matrix (100% indicates perfect clustering). **C)** Cluster stability plots showing that as the number of clusters increases past two, cluster stability decreases. **D)** PCA plot of the two predicted clusters, coloured by time-point, sized by infection status and shaped by cluster. **E)** PCA plot following kNN-smoothing on the expression matrix.

**Supplemental File 8. Sub-clustering**

The four comparisons shown here were created by manually selecting two and three clusters to examine any sub-clustering events not automatically detected. **A)** 3-hour cells - no sub-clustering evident. **B)** 6-hour cells - no apparent sub-clustering with two clusters; three clusters do display more of a separation (between blue and green), while infected and mock-infected cells cannot be distinguished. **C)** 12-hour cells - some separation evident with 3 clusters, but infection state is not distinguishable. **D)** Extracting only infected cells show a clear separation of 3 hour cells, but not 6 and 12 hours cells

**Supplemental File 9. Pseudotime analysis**

**A)** Pseudotime analysis of infected cells predicts three cell states. The minimum spanning tree (black line) is uninformative and not a true indication of an expected infection trajectory encompassing all cells from 3 to 12 hours. **B)** Each cell ordered throughout the predicted pseudotime and separated by cell state. **C)** Manually increasing the number of cell states to six appears to show a more realistic infection trajectory with a wider number of cells, in addition to showing start and end points correlating to 3− and 12-hour cell. **D)** When all cells are used, two cell states are predicted that enforce initial clustering. When the number of cell states is manually increased, smaller subsets appear providing a finer resolution.

**Supplemental File 10. Differentially expressed genes and enriched pathways**

**A)** Differentially expressed genes from infected and mock-infected cells at 6-hours and 12-hours. When comparing cells from cluster 1 against cluster 2, a more complex experimental design was needed that took into consideration the variety of underlying cells. **B)** Enriched pathways from Reactome using differentially expressed genes from **A**.

## Declarations

### Abbreviations

STI: Sexually transmitted infections
PID: Pelvic inflammatory disease
TFI: Tubal factor infertility
EBs: Elementary bodies
RBs: Reticulate bodies
scRNA-Seq: Single cell RNA sequencing
RLE: Relative log expression
DE: Differentially expressed
IF: Intermediate filament
MT: Metallothionein
FDR: False discovery rate
ROS: Reactive oxygen species
ncRNAs: Non-coding RNAs

### Consent for publication

Not applicable.

### Author contributions

RH analysed, interpreted and co-wrote the manuscript. JM assisted with analysis and interpretation. MH performed the chlamydial infections and scRNA-Seq laboratory methods. WH assisted with interpretation of the data and contributed to the manuscript. GM conceived the experiments, obtained the funding, oversaw the sequencing, data analysis and interpretation, and co-wrote the manuscript.

## Acknowledgements

Sequencing was performed at the Genome Resource Centre, Institute for Genome Sciences, University of Maryland School of Medicine. Data was analysed on the ARCLab high-performance computing cluster at UTS, with files hosted using the SpaceShuttle facility at Intersect Australia.

