## Supplemental Figures for "Early transcriptional landscapes of *Chlamydia trachomatis*-infected epithelial cells at single cell resolution"

Supplemental Figure 1

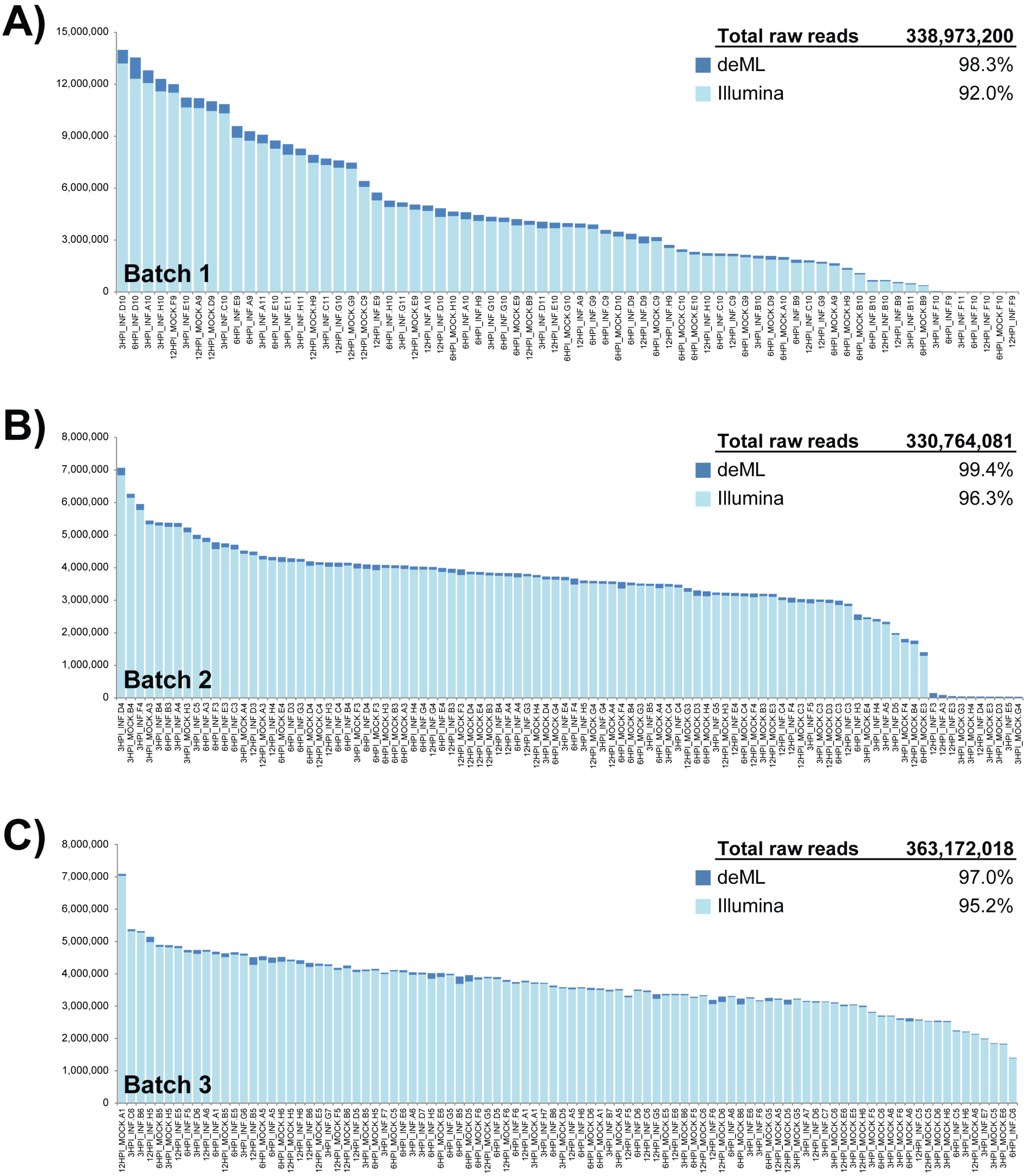

Supplemental Figure 2

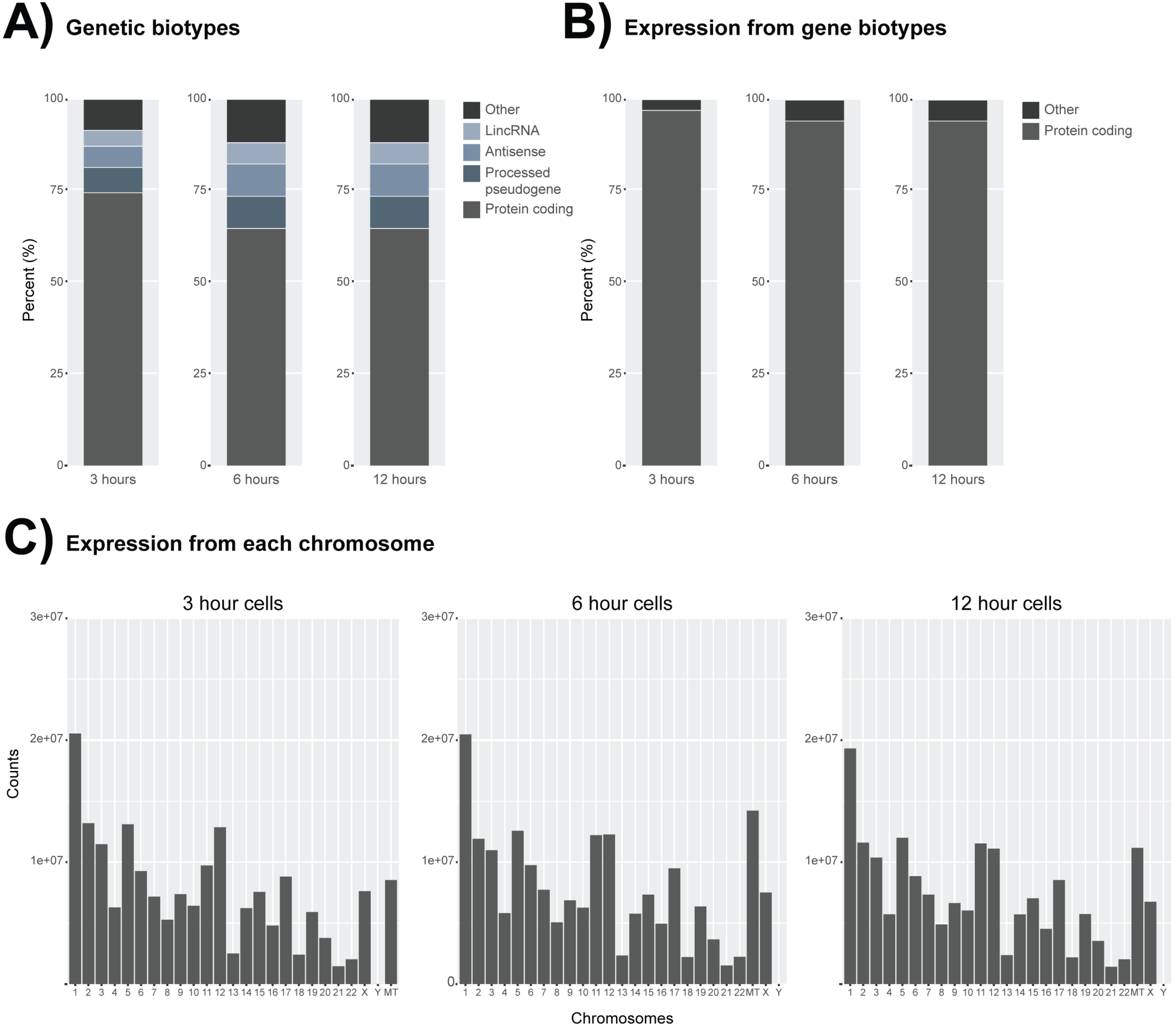

Supplemental Figure 3

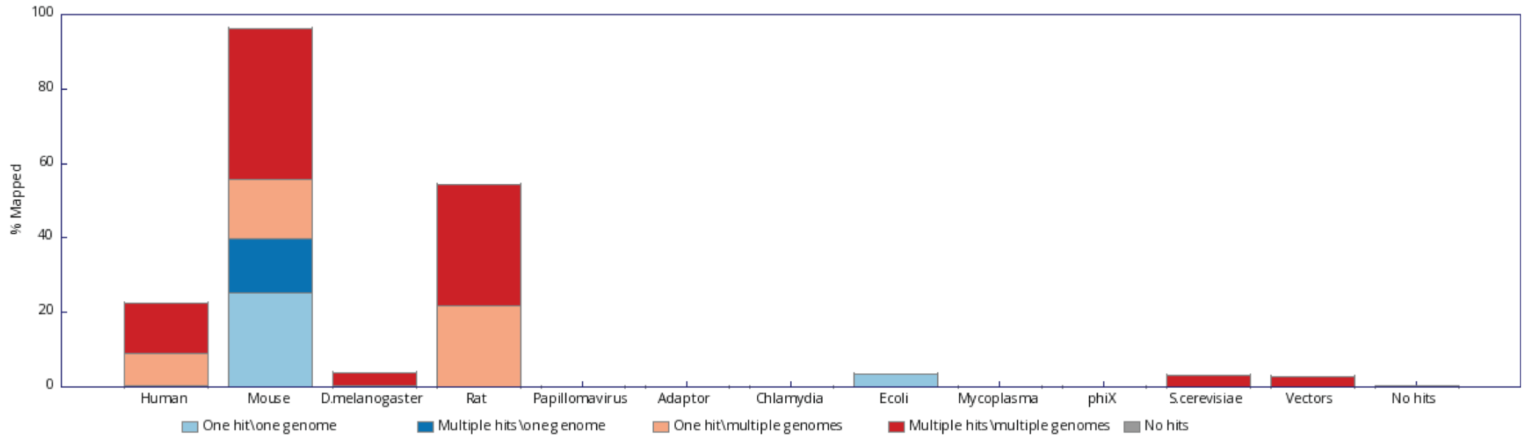


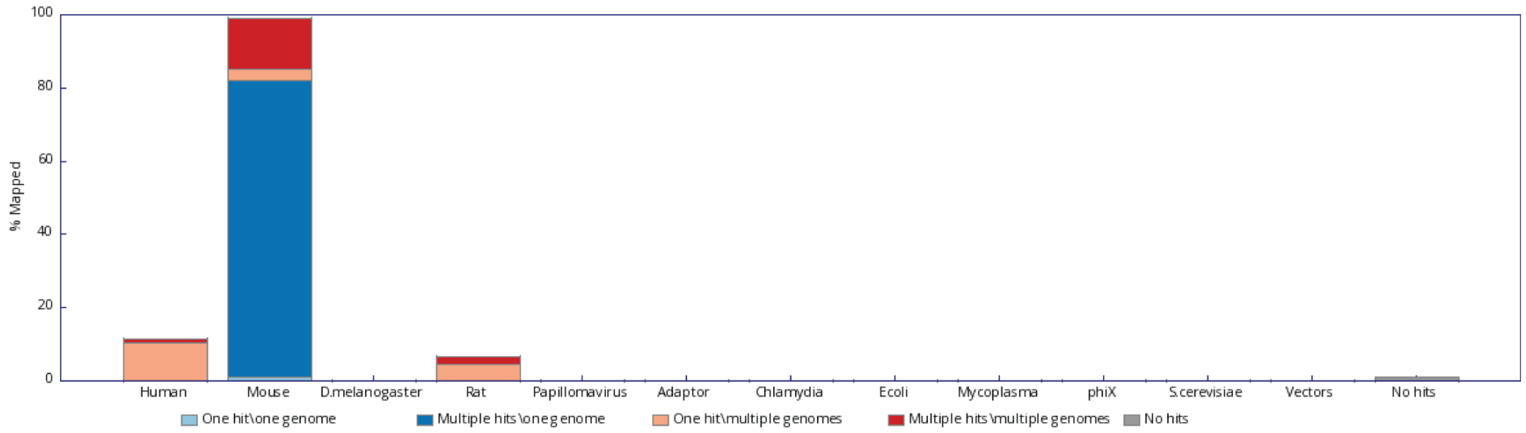

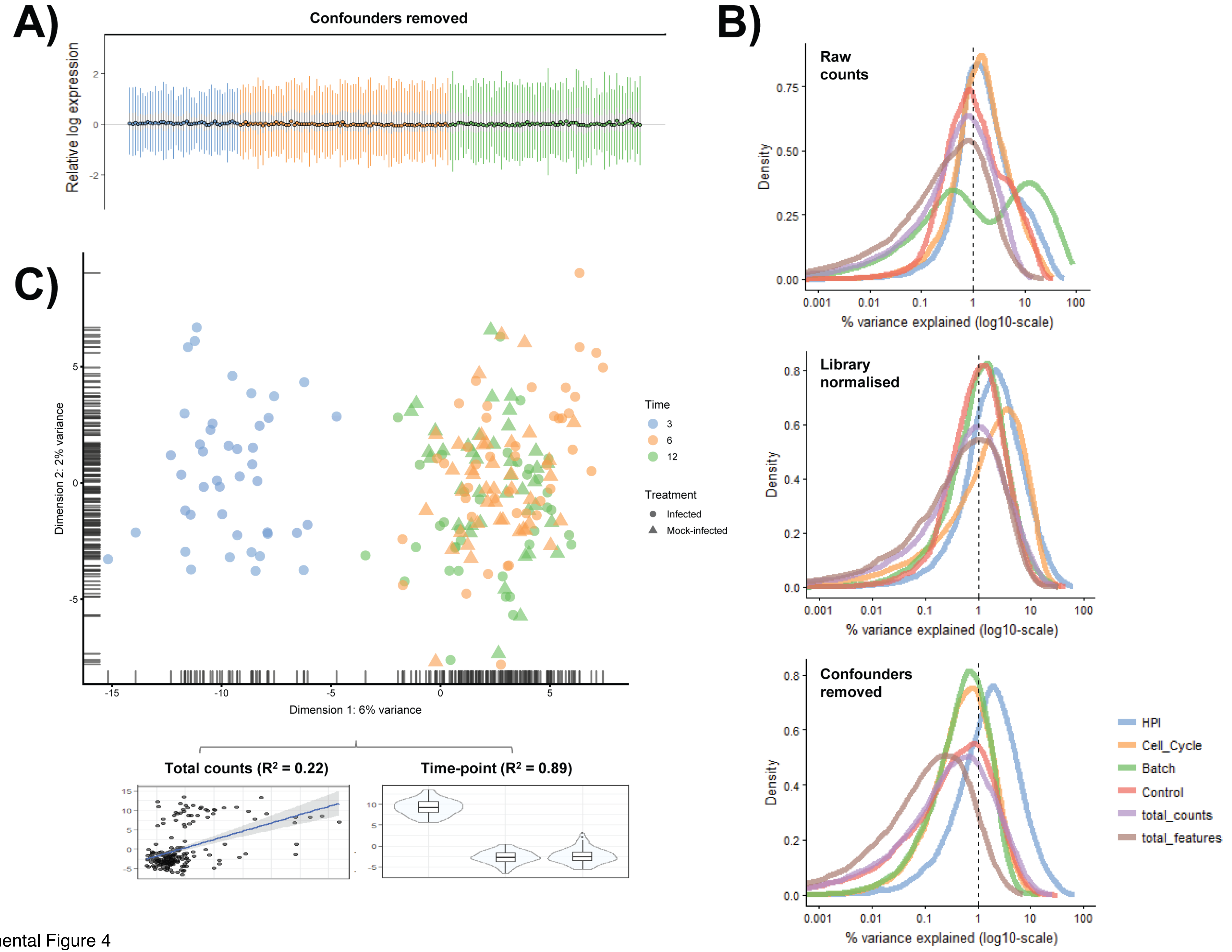


Supplemental Figure 4

Supplemental Figure 5

A)

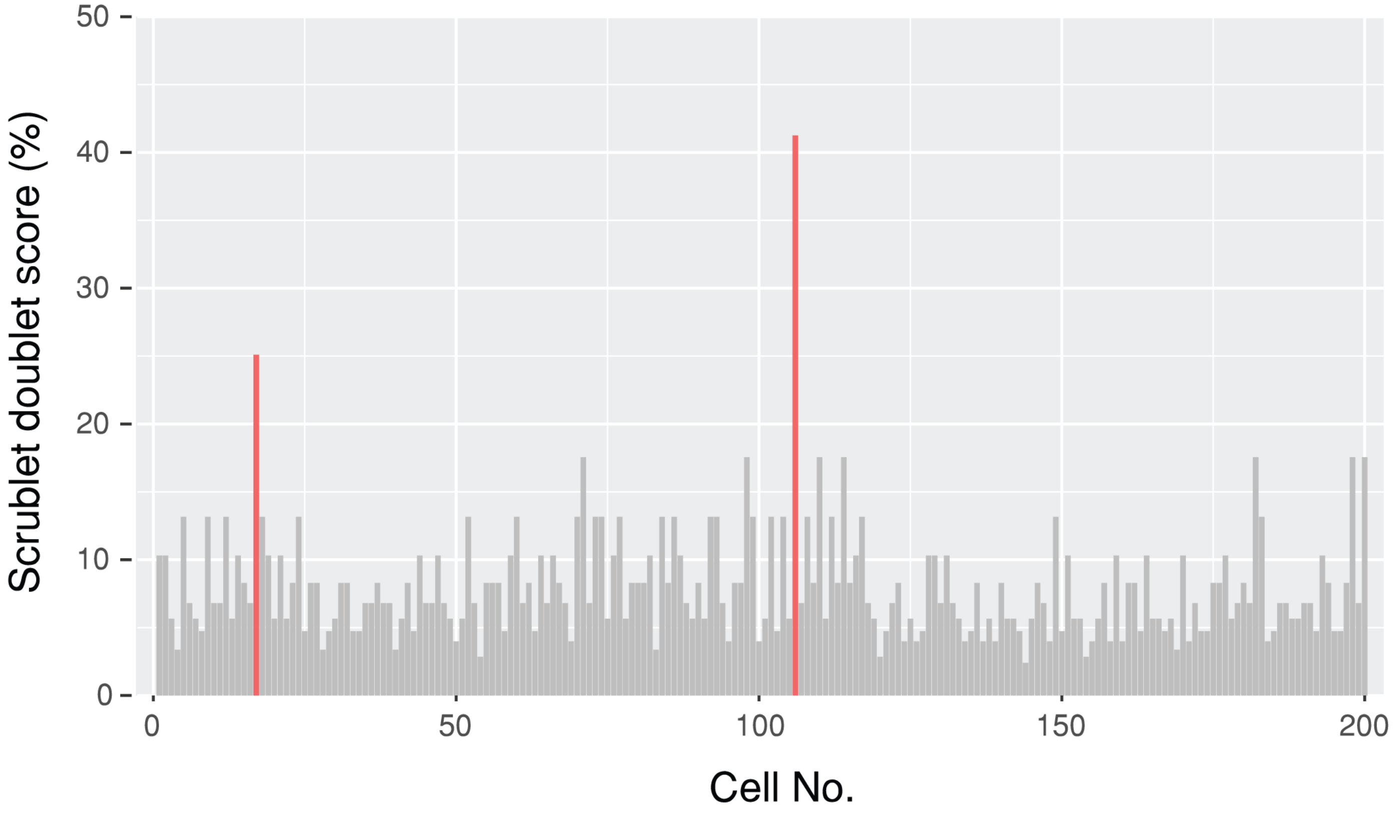


B)

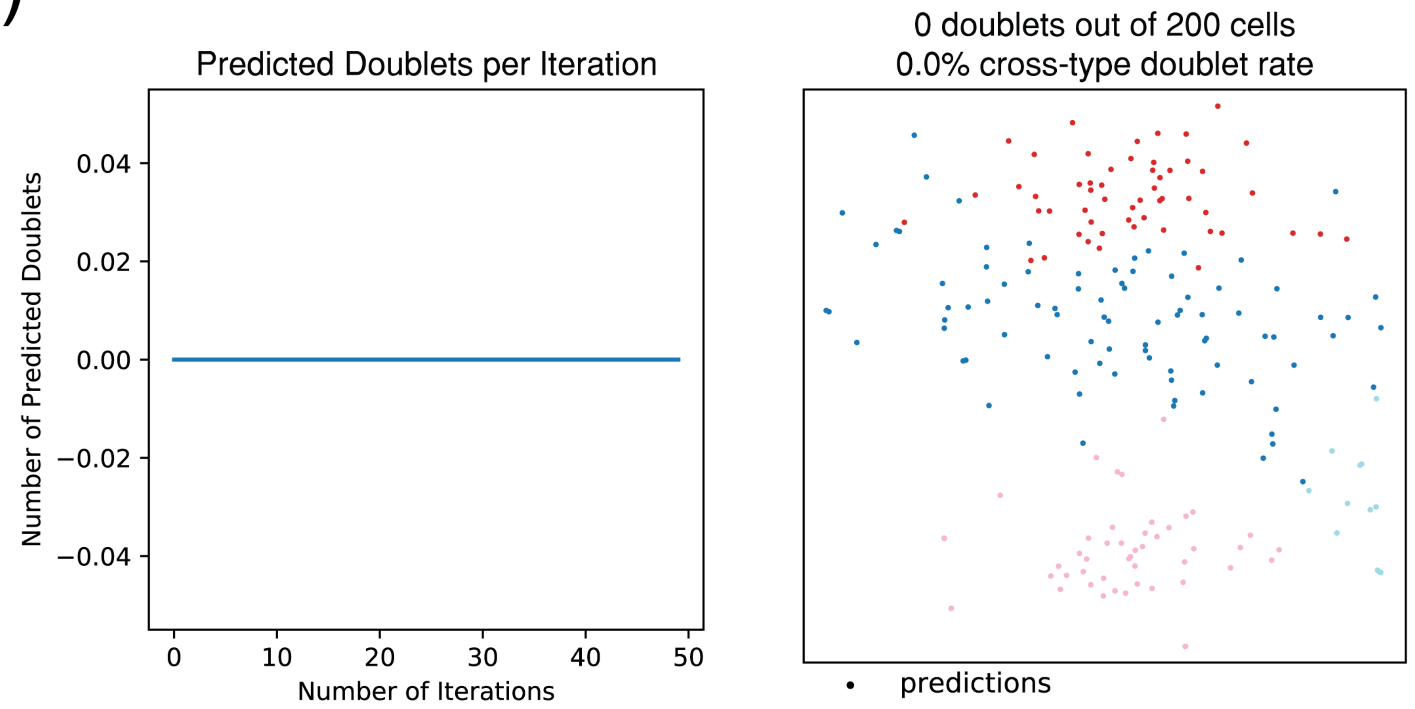

Supplemental Figure 6

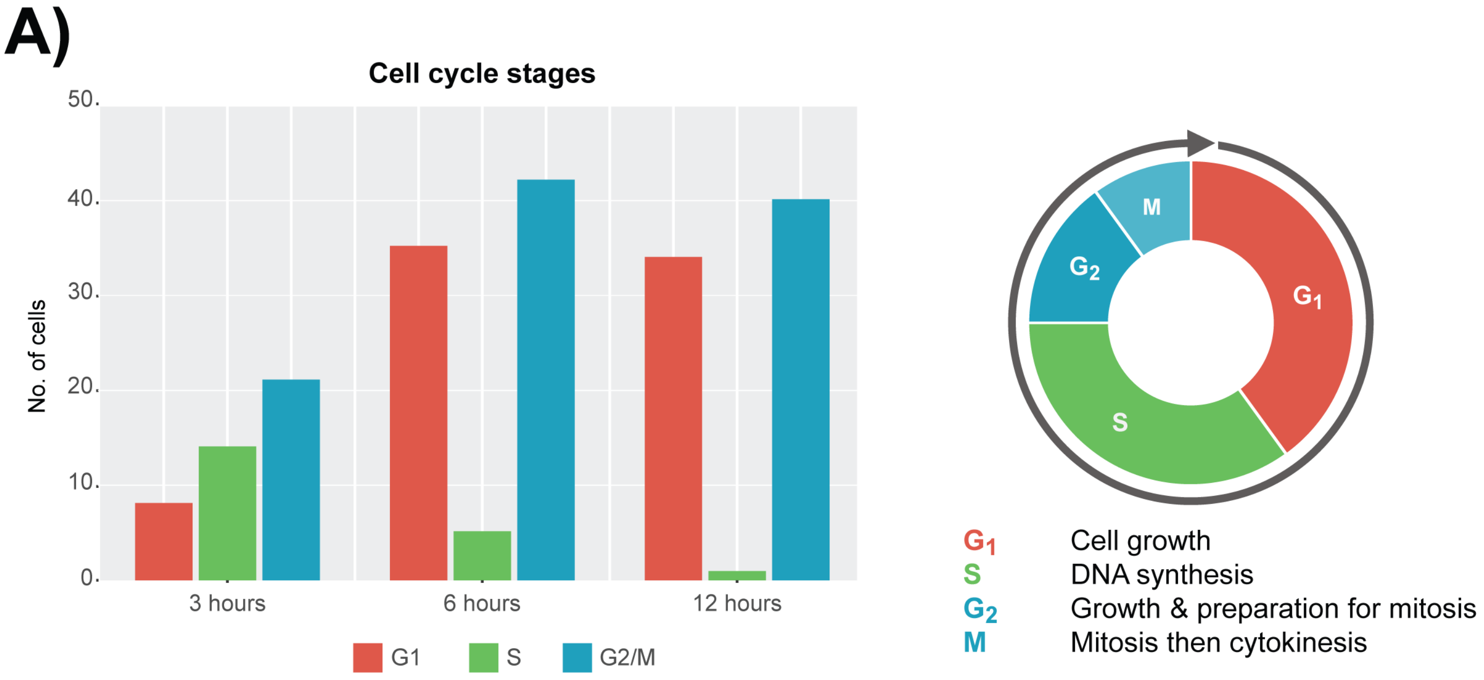


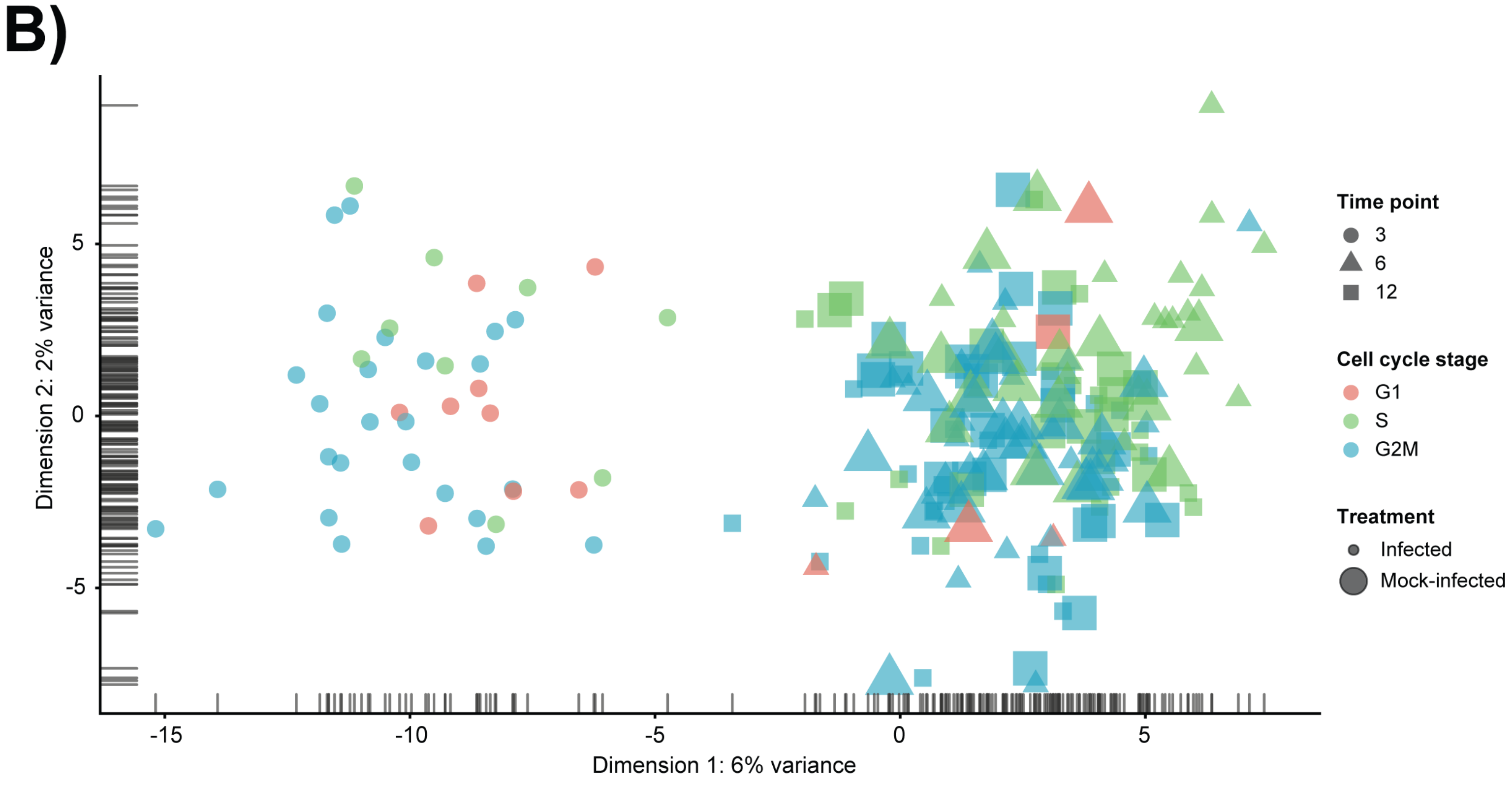

Supplemental Figure 7

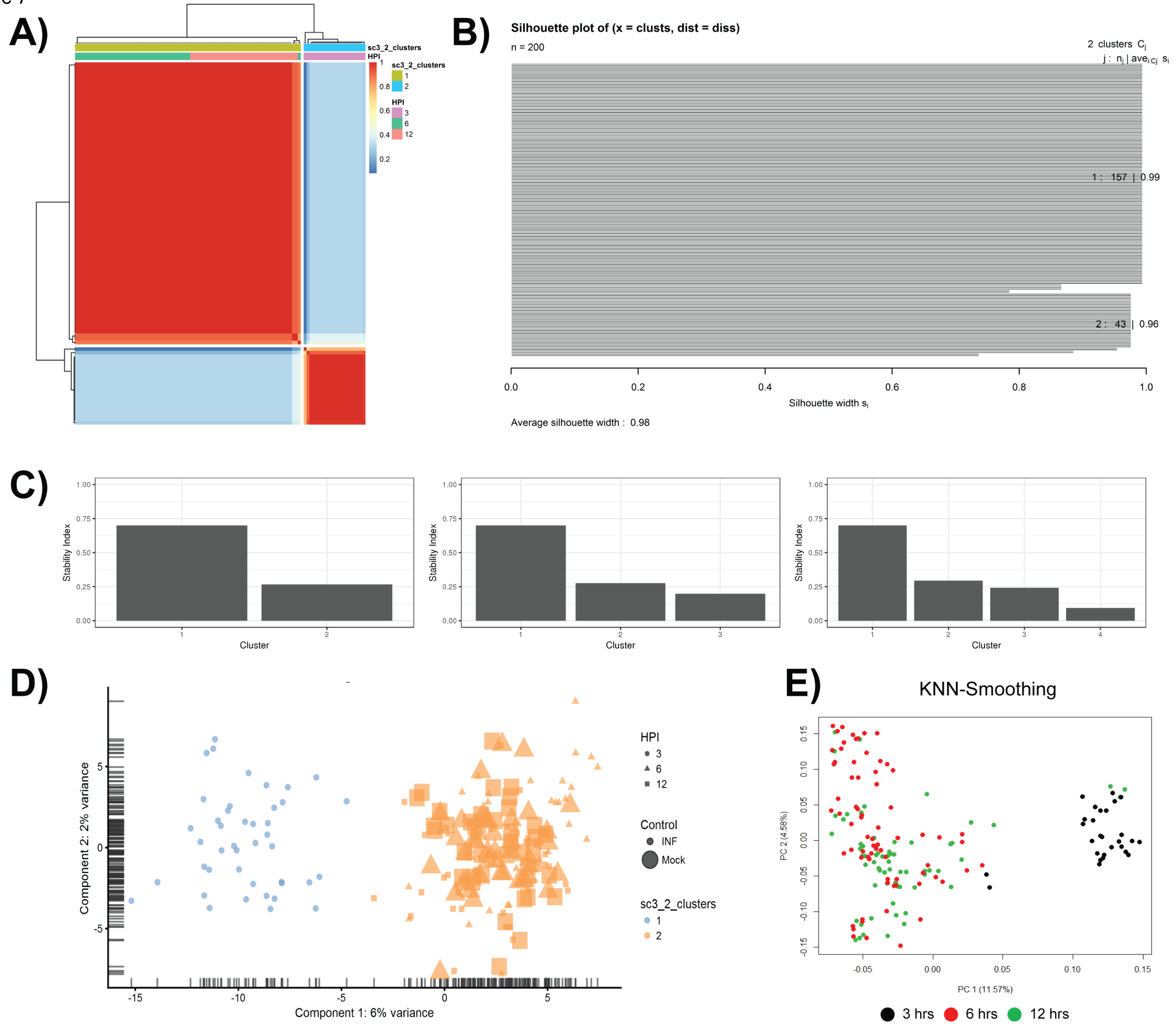

Supplemental Figure 8

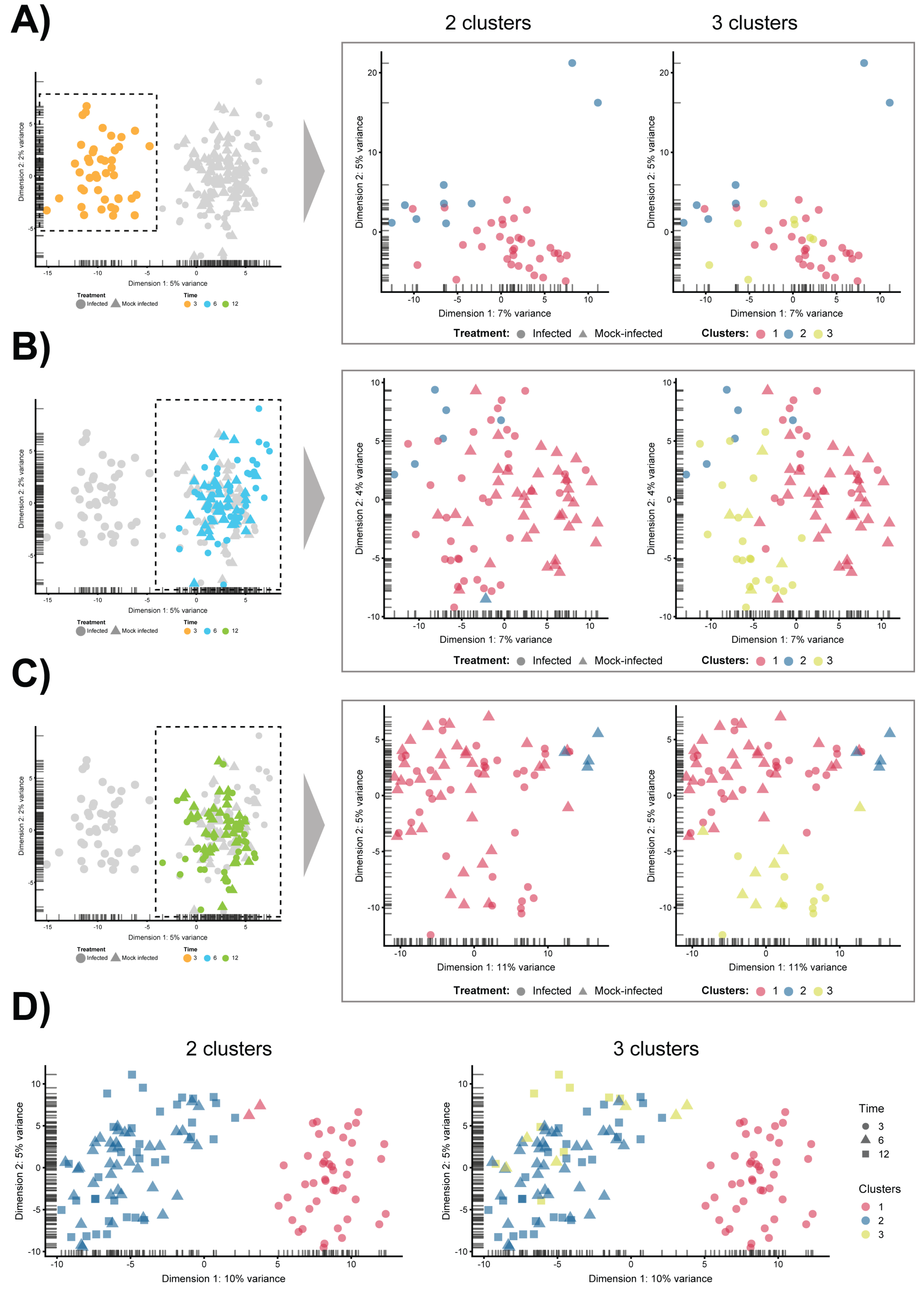

Supplemental Figure 9

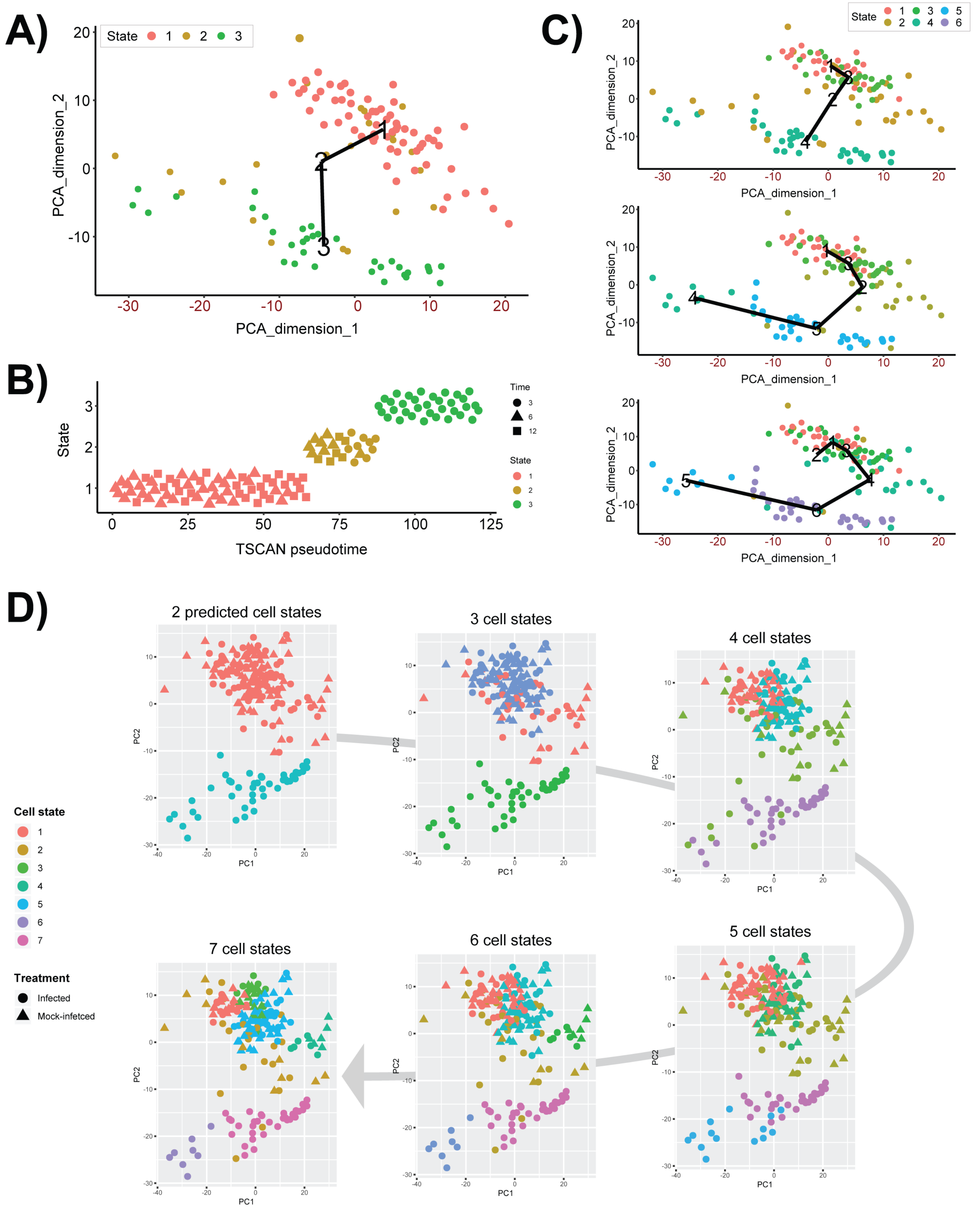

Supplemental Figure 10

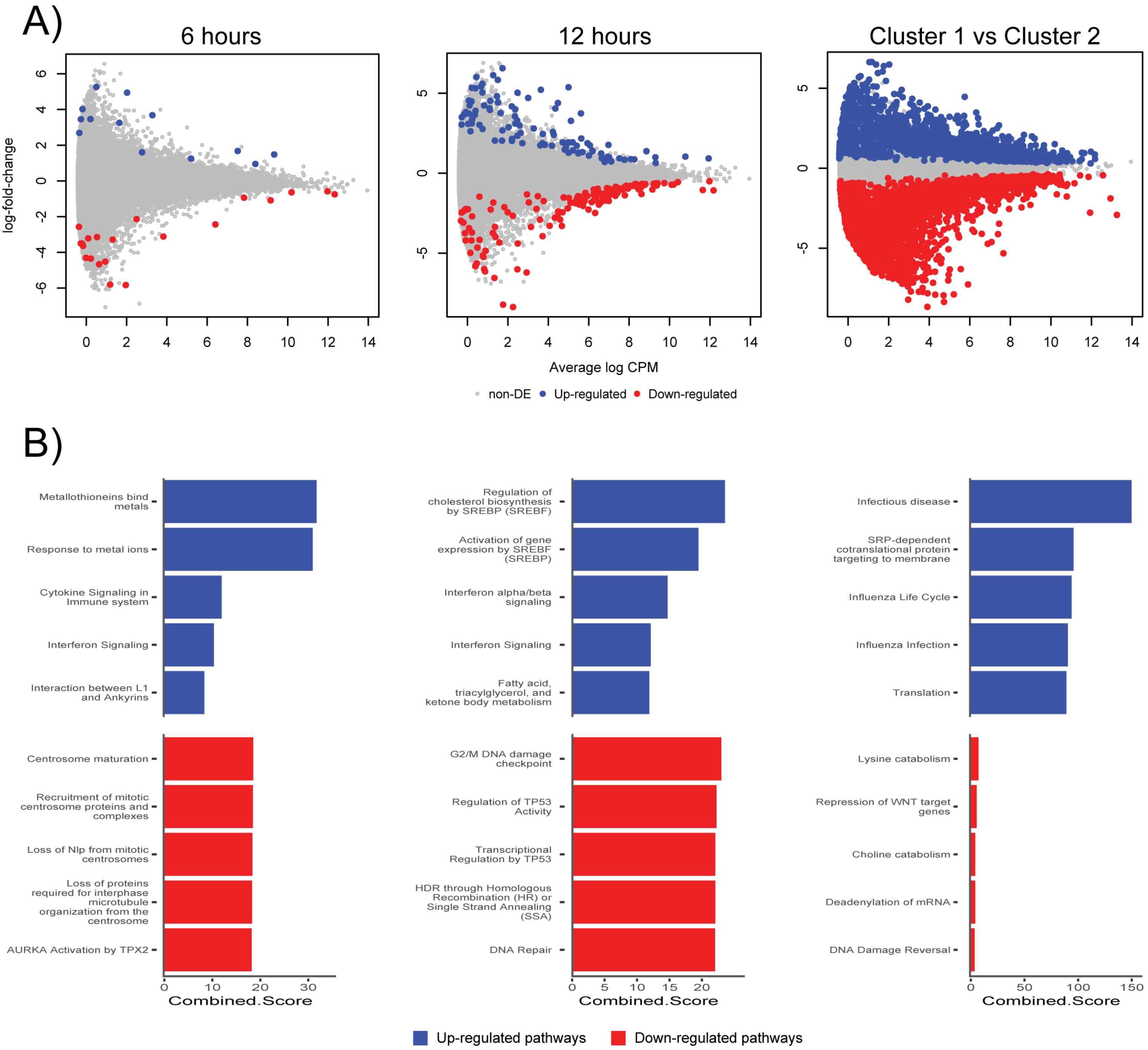
